## Supplementary material for "Caspar specifies primordial germ cell count and identity in *Drosophila melanogaster*": Suppl-FigMov2

### Slide 1
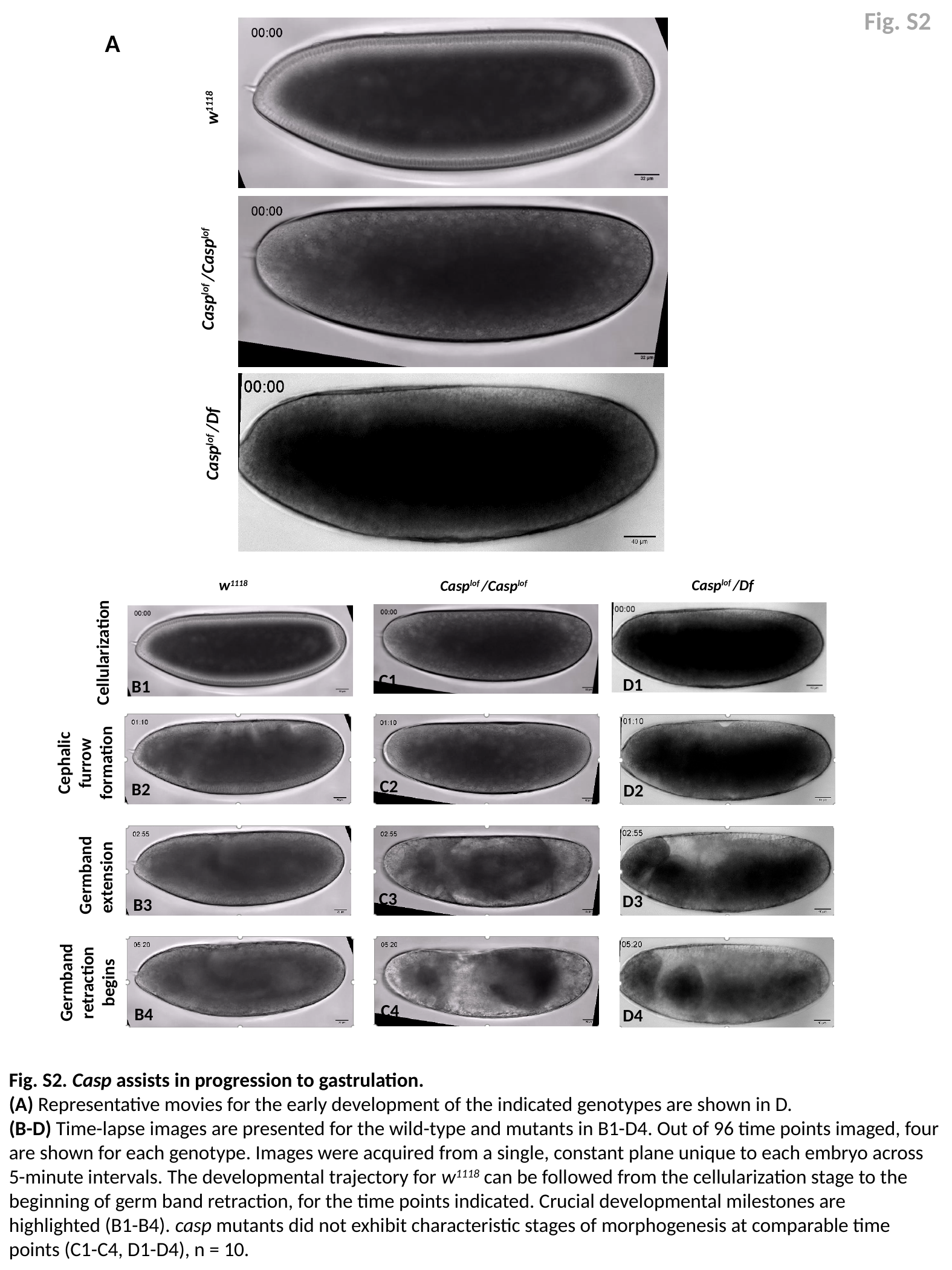

Fig. S2
A
w^1118^
Casp^lof^ /Casp^lof^
Casp^lof^ /Df
Casp^lof^ /Df
w^1118^
Casp^lof^ /Casp^lof^
Cellularization
C1
D1
B1
Cephalic furrow formation
C2
B2
D2
Germband extension
C3
D3
B3
Germband retraction begins
C4
B4
D4
Fig. S2. Casp assists in progression to gastrulation.
(A) Representative movies for the early development of the indicated genotypes are shown in D.
(B-D) Time-lapse images are presented for the wild-type and mutants in B1-D4. Out of 96 time points imaged, four are shown for each genotype. Images were acquired from a single, constant plane unique to each embryo across 5-minute intervals. The developmental trajectory for w^1118^ can be followed from the cellularization stage to the beginning of germ band retraction, for the time points indicated. Crucial developmental milestones are highlighted (B1-B4). casp mutants did not exhibit characteristic stages of morphogenesis at comparable time points (C1-C4, D1-D4), n = 10.
