## Supplementary material for "Caspar specifies primordial germ cell count and identity in *Drosophila melanogaster*": Suppl. Table 1

**Suppl. Table 1: Primers used for cloning casp constructs.**

| Name | Primer sequence (5'-3') | Use |
|---|---|---|
| *casp_F1* | *gtgccgttagcagcagtagc* | Forward primer for validating *casp$^{lof}$* lines. |
| *casp_F2* | *caactggtcacactgactagcc* | Forward primer for validating *casp$^{lof}$* lines. |
| *casp_R1* | *ccaatcgaacgaaacggcc* | Reverse primer for validating *casp$^{lof}$* lines. |
| *casp_R2* | *accggtagtatctggtgctg* | Reverse primer for validating *casp$^{lof}$* lines. |
| *casp_UAS_Ubx_F* | *ggacagtgagagctcgacagat* | Forward Primer for domain deletion mutants |
| *casp_Ubx_F* | *aaggcagagcaggacatggc* | Forward Primer for Casp$^{\Delta Ubx}$ mutants |
| *cmPUASPattB* | *ataggccactagtggatctgatgtacccat acgatgttccagattacgctggcggc* | Primer for adding the pUASPattB 5' homology arm to inserts |
| *pUASCasp_F* | *tgttccagattacgctggcggc* | Primer for amplifying the UAS_Casp inserts |
| *pUASCasp_R* | *accatgggtttaggtataatgttatcaagctc c* | Primer for amplifying the UAS_Casp inserts |
| *pUASddR* | *ttaggtataatgttatcaagctcctcacgtgg agccacttatctcatcatccg* | Reverse primer for amplifying the UAS_Casp_ΔUAS_ΔUbx insert |
| *pUASdUAS_F2* | *cggatgatgagataagtggctccacggaa acatgcgaaatgtttgaggagcag* | Forward primer for amplifying the fragment downstream of UAS domain |
| *pUASdUAS_R1* | *cgtggagccacttatctcatcatccg* | Reverse primer for amplifying the fragment upstream of UAS domain |
| *pUASdUBA_F* | *tgttccagattacgctggcggcccccatcct atcctggtgcc* | Forward primer for amplifying the UAS_Casp_ΔUBA insert |
| *pUASdUbx_R* | *ttaggtataatgttatcaagctcctcattcgg acggctcctgaggtag* | Reverse primer for amplifying the UAS_Casp_ΔUbx insert |
